# Machine Learning and Metabolic Model Guided CRISPRi Reveals a Central Role for Phosphoglycerate Mutase in *Chlamydia trachomatis* Persistence

**DOI:** 10.1101/2023.12.18.572198

**Authors:** Niaz Bahar Chowdhury, Nick Pokorzynski, Elizabeth A. Rucks, Scot P. Ouellette, Rey A. Carabeo, Rajib Saha

**Affiliations:** Chemical and Biomolecular Engineering, University of Nebraska-Lincoln, Lincoln, Nebraska, 68508, USA; Department of Pathology, Microbiology, and Immunology, University of Nebraska Medical Center, Omaha, Nebraska, 68198, USA

**Keywords:** *C. trachomatis*, persistence, nutrient starvation, global stress response, metabolic bottleneck

## Abstract

Upon nutrient starvation, *Chlamydia trachomatis* serovar L2 (CTL) shifts from its normal growth to a non-replicating form, termed persistence. It is unclear if persistence is an adaptive response or lack of it. To understand that transcriptomics data were collected for nutrient-sufficient and nutrient-starved CTL. Applying machine learning approaches on transcriptomics data revealed a global transcriptomic rewiring of CTL under stress conditions without having any global stress regulator. This indicated that CTL’s stress response is due to lack of an adaptive response mechanism. To investigate the impact of this on CTL metabolism, we reconstructed a genome-scale metabolic model of CTL (*i*CTL278) and contextualized it with the collected transcriptomics data. Using the metabolic bottleneck analysis on contextualized *i*CTL278, we observed phosphoglycerate mutase (*pgm)* regulates the entry of CTL to the persistence. Later, *pgm* was found to have the highest thermodynamics driving force and lowest enzymatic cost. Furthermore, CRISPRi-driven knockdown of *pgm* and tryptophan starvation experiments revealed the importance of this gene in inducing persistence. Hence, this work, for the first time, introduced thermodynamics and enzyme-cost as tools to gain deeper understanding on CTL persistence.

## Introduction

*Chlamydia trachomatis* serovar L2 (CTL) is a Gram-negative obligate intracellular human pathogen^1^ causing an estimated 1.7 million new genital tract infections annually^2^. This obligate intracellular pathogen replicates in a specialized membrane compartment, named inclusion, and uses different strategies to survive in the host intracellular environment^3^. During its development cycle, CTL can alternate between the extracellular infectious elementary body (EB) and the intracellular non-infectious reticulate body (RB)^4^. EBs enter epithelial cells and differentiate into RBs inside the inclusion. After several cycles of replication, RBs undergo secondary differentiation to create new EBs, which promote subsequent rounds of host cell infection^5^.

When RBs sense unfavorable growth environments, such as lack of essential nutrients or immune responses, it enters a reversible state of growth-arrest, termed persistence^6^. The RB in the persistent state exhibits a distinct morphological form, referred to as an aberrant reticulate body (ARB). ARBs are metabolically active with an enlarged and aberrant morphology^7^. Several studies identified stimuli that induce CTL persistence. Some of these stimuli are the addition of the cytokine interferon-γ^8^, tryptophan starvation^9^, iron starvation^10^, and infection with immune cells^11^. Despite these studies, the fundamental nature of persistence is still puzzling. One hypothesis is, CTL enters persistence as it lacks an adaptive response strategy^12–14^. An extension of this hypothesis is that persistence is effectively a substitute for the lack of a conventional global response regulator, when encountering nutritional stress. A recent study^15^ compared iron and tryptophan starvation to the nutrient-replete conditions. While there were significant transcriptomic differences between control and starvation conditions, it is still not clear if those changes are due to CTL’s lack of adaptive response on nutrient starvation or not. To gain a holistic understanding, it is pertinent to understand the metabolic landscape corresponding to the transcriptional changes. However, there is a dearth of experimental studies investigating the whole CTL metabolic landscape. Thus, a systems biology approach is required to understand the metabolism of CTL under different stress conditions.

To develop such a systems biology approach, genome-scale metabolic models (GSMs) have been widely used^16,17^. A GSM captures annotated metabolic reactions within a biological system and can predict reaction fluxes using flux balance analysis (FBA)^18^, flux variability analysis (FVA)^19^, and parsimonious FBA (pFBA)^20^. To date, GSMs developed for different pathogens were successful in predicting metabolic adaptations, functionalities, and properties of *Mycobacterium tuberculosis*^21^, *Acinetobacter baumannii*^22^, *Klebsiella pneumoniae*^23^, and *Helicobacter pylori*^24^. GSMs were also successful in predicting different virulence factors such as lipid A modifications of *Pseudomonas aeruginosa*^25^, and substrate utilization pattern of *Staphylococcus aureus*^26^. Moreover, GSMs successfully predicted novel drug targets for *Salmonella typhimurium*^27^, and *Campylobacter jejuni*^28^. Hence, to understand the metabolic landscape of CTL, GSM can be useful. There is a recently published GSM of *C. trachomatis*^29^. However, the biochemical network of the model was incomplete. To address this, we reconstructed a new GSM of CTL, *i*CTL278.

To study the nature of persistence, persistence-specific ‘omics’ data needed to be overlayed with the *i*CTL278. This process is called contextualization of the GSM. Without contextualization, GSM may predict unrealistic reaction fluxes^30^, erroneous cellular phenotype^31^, or inaccurate growth rate patterns^32^. Two different approaches are available for contextualization, switch (e.g., GIMME^33^, iMAT^34^, MADE^35^, and RIPTiDe^36^) or valve (e.g., E-Flux^37^ and PROM^38^, EXTREAM^39^) approach. While switch approaches display a binary nature, resulting in the active-or-inactive status of a reaction, valve approaches provide more flexibility by making the reaction flux proportional to the abundance of associated transcripts/proteins.

In this work, to perform systems-level investigations of CTL persistence, transcriptomics data were collected for nutrient-sufficient and nutrient-starved conditions. To gain further insights from these transcriptomics data, an unsupervised machine learning approach called K-means clustering algorithm was implemented resulting in the identification of four core components of the CTL transcriptome. This along with a statistical correlation study revealed a global transcriptomic rewiring of CTL under nutritional stress conditions. This, along with a lack of global stress regulator in CTL revealed lack adaptive response as the primary reason of CTL’s entry to persistence. To further understand how this global transcriptomic response impacts CTL metabolism, contextualized GSMs were reconstructed using the E-flux algorithm. Our recently developed Metabolic bottleneck analysis (MBA)^39^ identified phosphoglycerate mutase (*pgm*) regulated the entry of CTL into persistence, which had the highest thermodynamics driving cost and lowest enzymatic cost. Furthermore, we used CRISPRi to block transcription of *pgm* and subsequently performed starvation experiments. The outcome supported the prediction from MBA, thermodynamics driving force, and enzyme cost analysis. Additional systems-level investigation pin-pointed the cellular fitness of the persistence, which is to prime itself for a rapid growth upon availability of nutrient. Overall, this combined machine learning and metabolic model-guided study, for the first time, decoded CTL persistence through the lens of thermodynamic driving force and enzymatic cost and will work as a blueprint to investigate phenotypical and genotypical changes associated with other microbial infection.

## Results And Discussion

### *Chlamydia trachomatis* L2 genome-scale metabolic model development and analysis

To analyze the metabolism of CTL in different stress conditions, we needed to reconstruct a GSM of CTL. A draft GSM was reconstructed using the NCBI RefSeq genome annotation from the Kbase^40^.

After reconstructing the draft GSM, an extensive literature search was performed regarding the CTL-specific biomass composition. However, with information unavailable, a template Gram-negative biomass equation was used as the objective function of FBA. This draft model did not capture some known metabolic functionality of CTL. For example, CTL has an incomplete TCA cycle and must uptake malate^41^. However, the malate uptake reaction was missing in the model. Similarly, CTL can uptake glucose 6-phosphate from the host cell^41^. To account for these, we performed gap-filling using our previously developed tool OptFill^42^ and added transport reactions for both malate and glucose 6-phosphate to the model. CTL partially depends on the host cell for ATP and NAD+ uptake^41^. We also added transport reactions for CTL to uptake these metabolites when necessary. Overall, the curated model, *i*CTL278, consisted of 729 metabolites, 692 reactions, and 278 genes. We performed the MEMOTE testing^43^ for *i*CTL278 and it returned an impressive score of 94%. The model is fully mass balanced and stoichiometrically consistent. The full MEMOTE report can be found in the supplementary S1 Text. The MEMOTE score comparison between *i*CTL278 and the previously published CTL model is shown in Fig. S1A. In addition to performing MEMOTE, we also verified other known metabolic traits of CTL, such as the *i*CTL278 can uptake several nucleotides including GTP, CTP, and UTP to sustain biomass growth^44^. CTL is an auxotroph for most amino acids and can only biosynthesize alanine, aspartate, and glutamine using dicarboxylates from the TCA cycle^41^. The rest of the amino acids need to be acquired from the host cell. *i*CTL278 recapitulated these metabolic traits of amino acids. Overall, *i*CTL278 captured all the known metabolic characteristics of CTL. The model reconstruction process is shown in Fig. 1, the metabolic networks of the model are shown in Fig. 2A and Fig. 2B and the number of reactions in different pathways is shown in Fig. 2C.

**Fig. 1.**
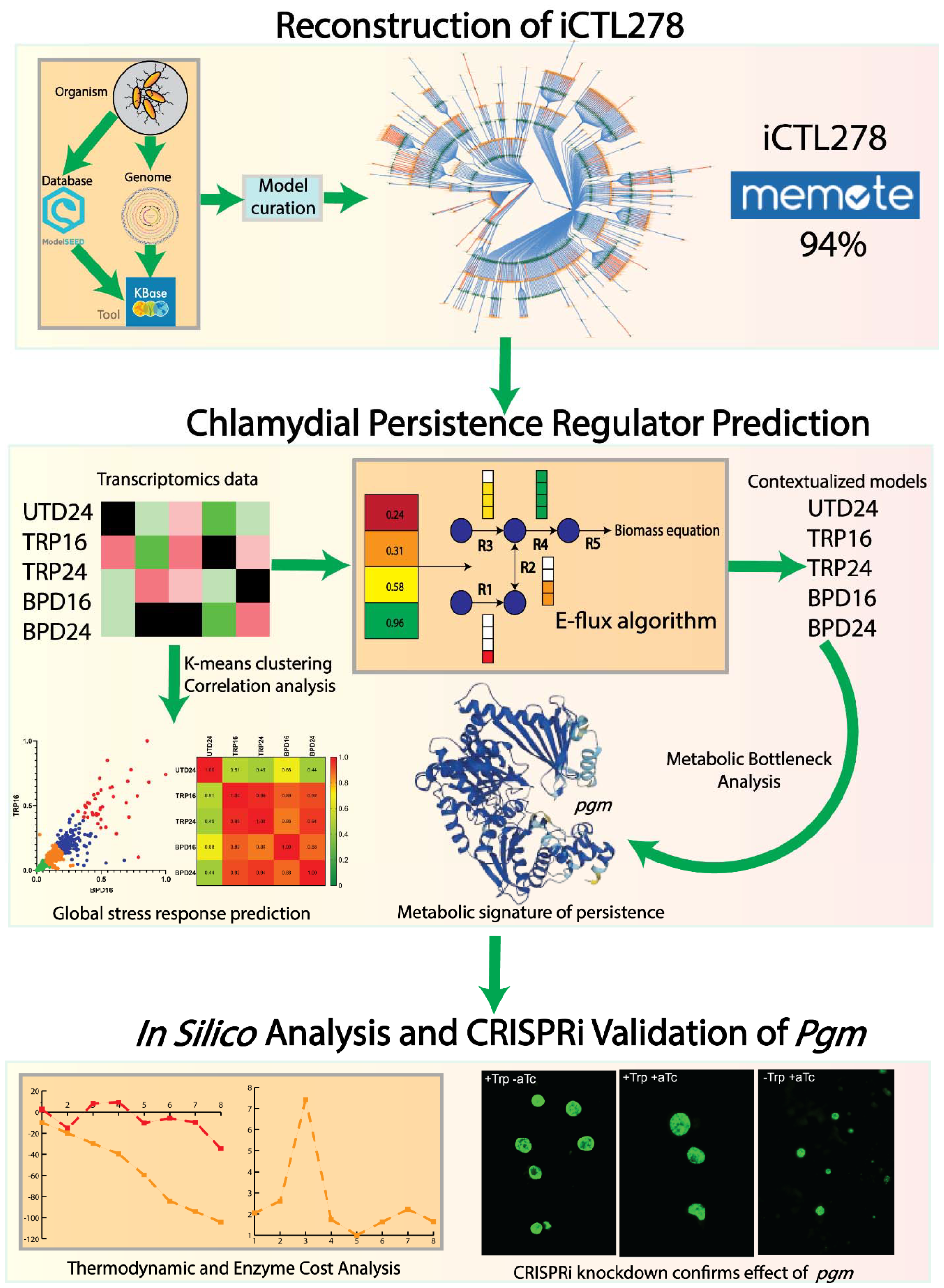
Overall process of the model reconstruction, refinement, and subsequent analysis/validation. Kbase, NCBI RefSeq, and Modelseed database was used to reconstruct the initial model. After standard model curations, a high-quality GSM was ready to use. Later, condition specific transcriptomics data were incorporated with GSM through E-flux algorithm, which resulted in five contextualized models. These models were used to find metabolic bottlenecks and persistence mechanism. K-mean clustering algorithm applied on the transcriptomics data revealed global stress response mechanism. Thermodynamics and enzyme cost analysis, along with *in vitro* experimentation reveal the regulatory role of *pgm* in CTL persistence.

**Fig. 2.**
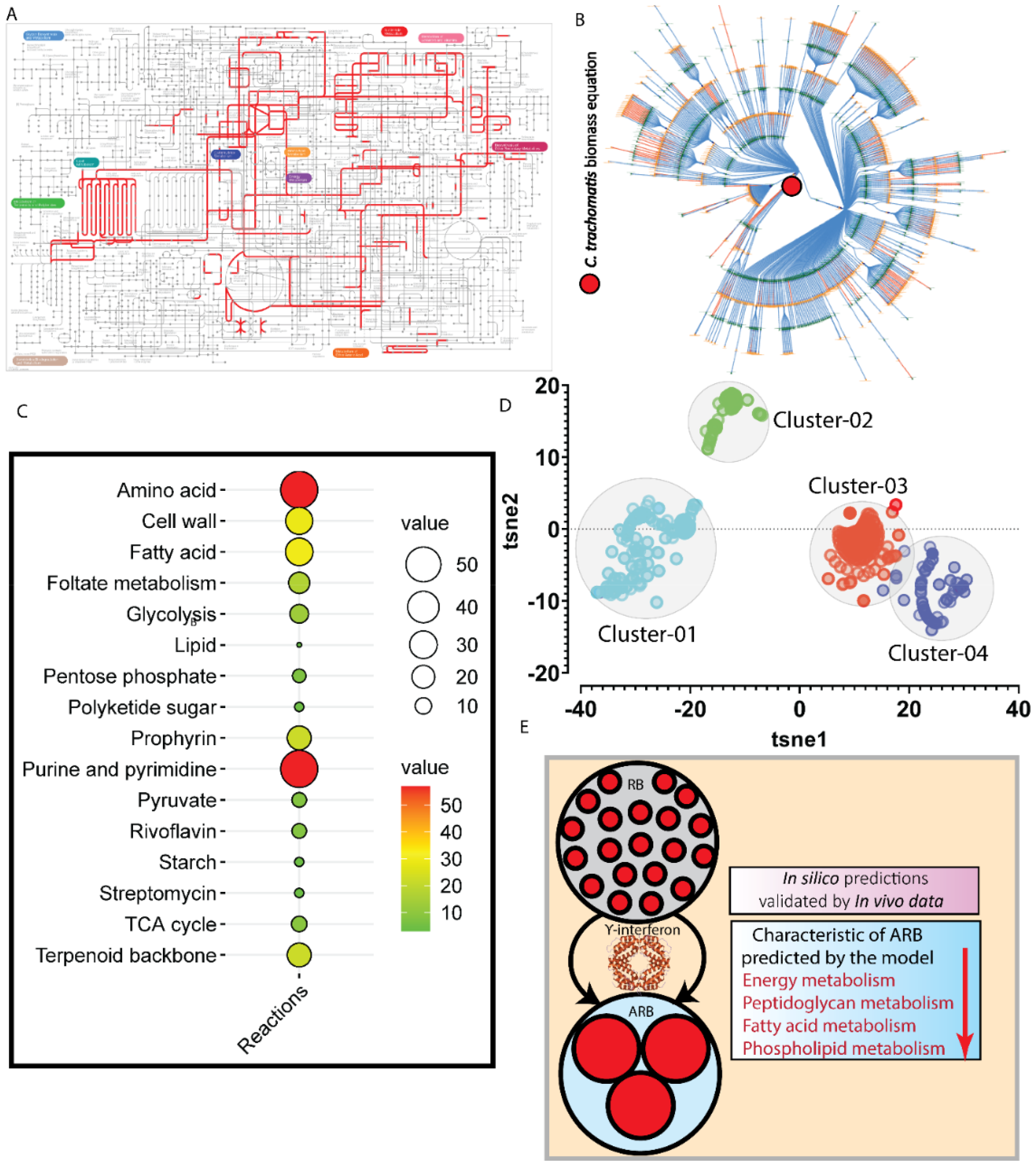
An overview of the metabolic network captured in the CTL genome-scale metabolic model and result of model validation. A) Visualization of the GSM metabolic network was performed using iPath3^45^. This metabolic network provides an idea regarding the limited metabolic capability of CTL; B) Metabolic network of CTL is centered around its biomass equation, network generate using fluxer^46^; C) A balloon plot representing number of reactions in different pathways in the *i*CTL278; D) tSNE plot of the flux sampling data obtained from *i*CTL278; E) Predictions for contextualized *i*CTL278 matched well with the experimental observations.

We also wanted to understand how the interconnectedness of the major pathways of *i*CTL278. Hence, we performed flux sampling of *i*CTL278, followed by the dimensionality reduction and clustering of those flux samples. The analysis yielded four different clusters (Fig. 2 D). Cluster-01 reactions are mostly from glycolysis and pentose phosphate pathways. In cluster-02, most of the reactions are from the ubiquinone and terpenoid pathways. Cluster-03 consists of primarily pyrimidine and fatty acid pathways. Finally, cluster-04 contains reactions mostly from purine and riboflavin pathways. From these clustering, we now have a better understanding how metabolic pathways in CTL are connected to one another in a higher dimensional space. Further details on the clusters can be found in supplementary S1 Table.

### Model Validation using experimental proteomics data

Although the MEMOTE testing of *i*CTL278 returned an impressive score of 94%, it was still necessary to determine if *i*CTL278 could capture the known phenotypes of CTL upon contextualization. In a previous study^37^, a valve-based approach, E-flux algorithm that connects reaction activities linearly with the gene expression levels accurately predicted different cellular phenotypes regarding fatty acid biosynthesis of *M. tuberculosis*, a well known pathogenic bacterium. Therefore, a previously published study on quantitative protein profiling of interferon-γ-treated CTL^8^ was used to reconstruct two contextualized models using the E-flux algorithm^37^, reticulate body (RB) and aberrant reticulate body (ARB) models. The latter was under stress induced by interferon-γ.

When flux distribution was calculated for both RB and ARB, similar to previous studies, flux through energy metabolism was reduced in the ARB compared to RB^8^. As CTL enters persistence, cell division is arrested, and metabolism is reduced to a minimum level such that only essential cellular functions are maintained. As GSMs are unable to directly calculate the concentration of different metabolites involved in a biochemical network, we used the flux-sum analysis (FSA) approach to predict the metabolic pool size of ATP in both RB and ARB for further verifications, and found a lower ATP pool size in the ARB condition. The details of flux-sum analysis can be found in our previous work^32^ and also in the materials and methods section. Thus, the reduced energy metabolism in ARB compared to the RB, a distinguishing feature of the persistence state, was successfully captured by the *i*CTL278.

As a further verification, we explored the peptidoglycan biosynthesis pathway of CTL. Peptidoglycan is essential cell division in CTB but, unusually, is not a component of the cell wall. During stress conditions, CTL arrests its cell division^6^. As cell division is arrested in the ARB, the model predicted reduced peptidoglycan biosynthesis in ARB compared to RB, which is consistent with a previous transcriptomic study^47^. CTL relies on *de novo* fatty acid and phospholipid biosynthesis to produce its membrane phospholipids, which are also crucial for cell division^48^. When fatty acid biosynthesis reactions in both RBs and ARBs were compared, we found reduced activity in the fatty acid biosynthesis in ARBs compared to RBs, which reiterates what has previously been reported^49^. This was expected given the predicted reduced requirement for fatty acid biosynthesis in the absence of bacterial replication.

Overall, *i*CTL278 recapitulates known phenotypes of ARBs and RBs upon contextualization. Therefore, *i*CTL278 is well suited to further contextualize and analyze the metabolism of CTL under different stress conditions. The summary of the validation result is shown in Fig. 2 E.

### Effects of nutrient starvation on CTL growth and development

Our understanding of the molecular underpinnings behind the persistence of CTL is still limited. Despite employing numerous methods to induce persistence, the result consistently shows similar ARB morphology, as well as disruptions in the transcriptome and proteome. Thus, severe alterations to transcriptional and translational profiles may cause aberrant growth.

We assayed several distinguishing features of CTL persistence across the different treatment conditions, i.e., untreated (UTD24), 16 hours of iron (BPD16) and tryptophan starvation (TRP16), and 24 hours of iron (BPD24) or tryptophan starvation (TRP24). One of those was the morphology of CTL, which was monitored by immunofluorescent confocal microscopy. Here we confirmed that, in contrast to the UTD24, all starvation strategies yielded smaller inclusions (Fig. 3 A), which was associated with lower yields of inclusion-forming units, indicating severely delayed development that compromised the regeneration of infectious particles. We observed that BPD treatments resulted in aberrantly enlarged organisms. TRP inclusions showed smaller inclusions, indicated growth defect. We also analyzed genome copy number across all stress conditions. We observed significantly reduced genome equivalents for all stress conditions compared to UTD24. Unsurprisingly, genome copy numbers of chlamydial late-stage transcriptional regulator, *euo*, between BPD and TRP were similar (Fig. 3 B). With regards to generation of infectious progeny, we observed a much more dramatic effect of BPD treatment than TRP starvation, indicating that iron starvation is a much more potent trigger of CTL developmental arrest relative to tryptophan depletion (Fig. 3 C).

**Fig. 3.**
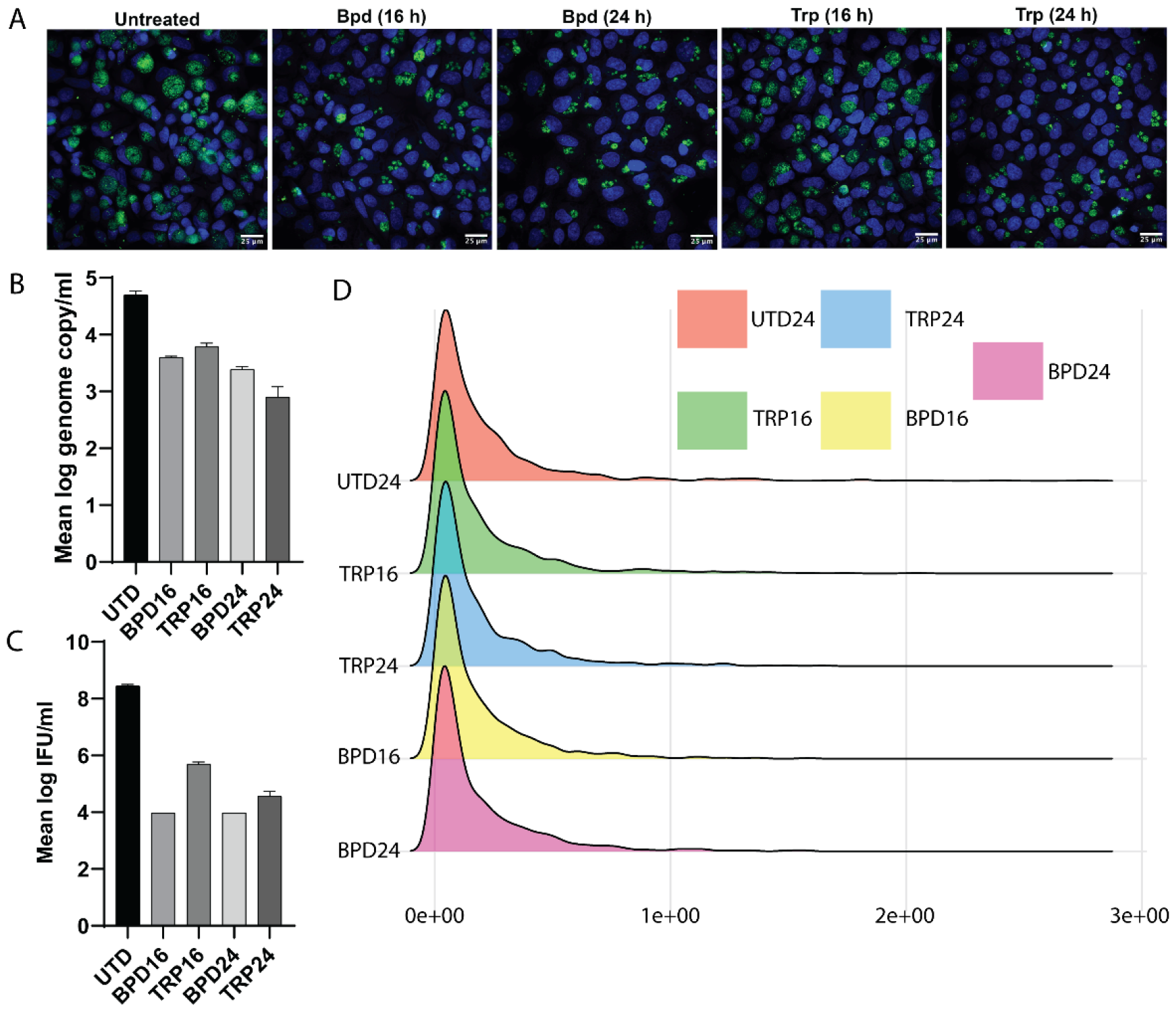
Morphology, genome-copy number, and infectious progeny data for different stage of CTL tryptophan and iron starvation. A) CTL exhibits duration-dependent sensitivity to iron limitation and tryptophan (trp) starvation. Infected cells were starved for iron by treatment with the chelator 2,2-bipyridyl (Bpd) starting at either the time of infection or at 8 h post-infection. Inclusions as indicated by staining for the chlamydial major outer membrane protein (MOMP) were allowed to develop for a total of 24 h. Tryptophan starvation was started at the same time points as above, with indicated durations. Note the more significant delay in inclusion development by Chlamydia starved for 24 h. B) Genome copy number data of *euo* gene for UTD24, BPD16, TRP16, BPD24, TRP24. C) Infectious progeny data for UTD24, BPD16, TRP16, BPD24, TRP24 D) Distribution of gene expression values.

To further analyze the stress conditions of CTL, RNA sequencing was performed for each of the stress conditions, including an untreated control^15^. Many genes fall in the low expression level and gene expression distribution also skewed heavily toward the low expression range (Fig. 3 D). Subsequently, a pair-wise comparison of profiles was conducted to determine the level of similarities of transcriptomes across different stress conditions. Using scatter plots, we were able to visualize correlations between pairs of transcriptomics data. From comparisons of TRP24-UTD24 (Fig 4. A), and BPD24-UTD24 (Fig 4. B), we found a weak correlation for each case (Fig 4. D). However, TRP24-BPD24 (Fig 4. C) showed a strong correlation (Fig 4. D). This pointed to CTL may having a common global transcriptomic response to different stress conditions. To reveal more about the pattern of global transcriptomic response, we implemented an unsupervised machine learning technique called K-mean clustering on TRP24-UTD24, BPD24-UTD24, and TRP24-BPD24. K-mean clustering identified four distinct clusters for each plot. Details of each cluster can be accessed in the supplementary S2 Table.

**Fig. 4.**
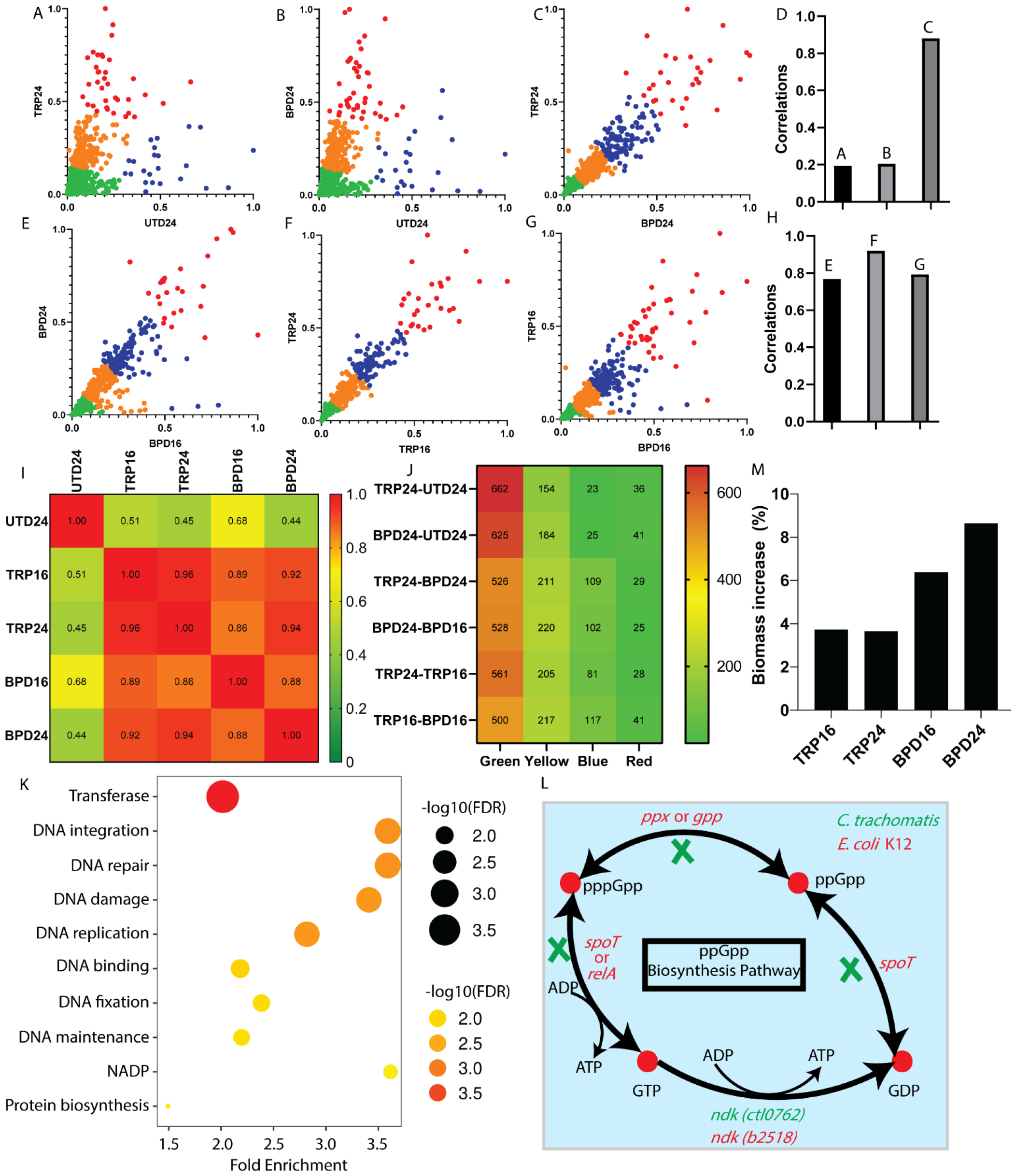
Pairwise K-mean clustering plots for different stress conditions. A) Scatter plot of 24 hour of tryptophan starved (TRP24) and untreated (UTD24) CT; B) Scatter plot of 24 hour of iron starved (BPD24) and untreated (UTD24) CT; C) Scatter plot of 24 hour of tryptophan starved (TRP24) and 24 hour of iron starved (BPD24); D) Correlation of different scatterplots. These scatterplots and correlation between different stress conditions reveal CTL may behave similarly under different stress conditions; E) Scatter plot of 24 hour of iron starved (BPD24) and 16 hour of iron starved (BPD16) CTL; F) Scatter plot of 24 hour of tryptophan starved (TRP24) and 16 hour of tryptophan starved (TRP16) CT; G) Scatter plot of 16 hour of tryptophan starved (TRP16) and 16 hour of iron starved (BPD16); H) Correlation of different scatterplots. These scatterplots and correlation between different stress conditions reveals, CTL may have a global transcriptomics response; I) Pearson correlation matrix among different conditions. The correlation was calculated using two-tail test with 95% confidence interval; J) Heat map for number of reactions in each cluster for different conditions; K) Gene ontology enrichment chart for common genes in green cluster across all conditions; L) Hypothetical ppGpp pathway in CTL; M) Increased biomass growths for stress condition after introducing ppGpp pathway in CTL.

Genes were clustered based on function, with clusters in green indicating metabolism-related genes; yellow clusters consisting of cell signaling-related genes, and red and blue clusters indicating transcription- and translation-related genes, respectively. From Fig. 4 (A, C), it is evident that most of the genes in green clusters remained similarly expressed. However, genes in the yellow and red cluster were moderately or highly upregulated in the TRP24/BPD24 compared to the UTD24 (Fig 4 A and B). In contrast, genes in the blue clusters were upregulated in the UTD24 compared to the TRP24/BPD24 (Fig 4 A and B).

As green clusters are tightly clustered across all the conditions (Fig 4. A, C), these can be called “core” components, as identified previously^15^. In contrast, blue, yellow, and red clusters were upregulated/downregulated across all conditions (Fig 4. A, C) and can be labeled as stress-specific “accessory” components^15^. Some of the known tryptophan utilization genes such as *trpR, trpC, and trpS* fell in the yellow clusters and were thus moderately upregulated in TRP24 (Fig 4. A). Several studies predicted similar upregulation of tryptophan utilization genes under tryptophan starvation^14^. *In vivo*, the amount of tryptophan available is often depleted by the pro-inflammatory cytokine interferon-γ through the transcriptional upregulation by the *ido1* gene, encoding the enzyme IDO that catabolizes tryptophan to kynurenine, which cannot be used by CTL. However the microbiome in the relatively hospitable niche of the lower genital tract may provide tryptophan or more likely indole, which CTL can use for tryptophan biosynthesis via the salvage pathway^50^. Indeed, several studies have shown that under tryptophan starvation, CTL can uptake indole supplemented in the media for conversion to tryptophan^50,51^.

Several stress-related genes, such as *ahpC* and *euo* were also upregulated in the TRP24 compared to the UTD24 (Fig 4. A). Between these two genes, *ahpC* was significantly upregulated and fell in the red cluster, while *euo* was moderately upregulated and included in the yellow cluster. *AhpC* is a thio-specific antioxidant peroxidase gene and is involved in the redox homeostasis of CTL. A similar severe upregulation of *ahpC* is also observed in CTL exposed to interferon-γ stress^47^. For *Chlamydia pneumoniae*, it was reported that severe upregulation of *ahpC* would protect the bacteria against cytokine-induced reactive nitrogen intermediates, thus allowing *C. pneumoniae* to cause long-term infection^52^. For *euo*, encoding a DNA-binding protein, a moderate upregulation was observed in CTL under interferon-γ induced stress. *Euo* was predicted to be a negative regulator of CTL genes involved in RB-to-EB differentiation^53^. *IhfA* is a cell division gene of CTL and was downregulated in the TRP24, and thus, possibly contributing to the lack of cell division observed in TRP24-treated samples (Fig 4. A).

In persistence, glycolysis mostly supports the increased production of starch and carbohydrate, thus CTL mostly depending on the host cell for energy^41^. As a result, reduced expression of TCA cycle genes, such as *sucA, sucB, sucC, and sucD* was expected, which fell in the bottom left portion of the green cluster. A similar overall result was observed for BPD24-UTD24 (Fig 4. B).

In the TRP24-BPD24 (Fig 4. C) scatterplot, the key difference is, that the stress-related accessory genes expressed more in chorus with the core genes, thus establishing a strong relation between two different stress conditions. Overall, the K-mean clustering predicted clusters whose functionality matched closely with the literature. Thus, this clustering analysis will further serve as a “genome-wide library” to identify core and stress-related genes of CTL.

To gather more insights into the presence of a global stress response in CTL, we plotted other stress conditions, such as BPD24-BPD16 (Fig 4. E), TRP24-TRP16 (Fig 4. F), and TRP16-BPD16 (Fig 4. G). Interestingly we noticed a very strong correlation in all the cases (Fig 4. H), supporting the proposed existence of global stress response of CTL. A Pearson correlation heat map (Fig 4. J) of normalized read counts of all genes among different conditions also indicated strong correlation between all the stress conditions. Number of reactions in each cluster for different conditions is shown in Fig. 4 I. Common genes in green, yellow, blue, and red clusters were identified using Venn diagrams (Fig. S1B, Fig. S1C, Fig. S1D, and Fig. S1E respectively). Gene ontology enrichment analysis for the common genes across green clusters is shown in Fig. 4 K. The same analysis for the yellow cluster (Fig. S2A), and the red clusters (Fig. S2B) are provided in the supplementary information. Since the blue cluster has only one common gene (*ctl0256*) we could not perform the gene ontology enrichment analysis.

Notably, the strong correlation between transcriptomes associated with two distinct stresses is not unique to CTL. Analysis of publicly available transcriptomes from differently stressed *Mycobacterium tuberculosis* and *Escherichia coli* revealed a similar correlation. We collected the transcriptional profile of *M. tuberculosis* exposed to *in vitro* lysosomal stress for 24 hours and 48 hours^54^. From the scatterplot (Fig. S3A), we noticed a very high correlation (Fig. S3D). We also collected the transcriptional profile of *M. tuberculosis* exposed in zinc limited medium and zinc replete medium^55^. Similar to the previous case, the scatterplot (Fig. S3B) indicated a very high correlation (Fig. S3D). However, when we scatterplot (Fig. S3C) *M. tuberculosis* transcription data for lysosomal stress of 24 hours and zinc limited medium, it showed a very low correlation (Fig. S3D). Thus, *M. tuberculosis* response to divergent stresses is customized to each stress. However, the transcriptional response for a single type of stress in *M. tuberculosis* progresses as a function of severity (i.e. duration) of the stress.

Unlike *M. tuberculosis, E. coli* showed a global stress response mechanism. Transcriptomics profile of 50 minutes of cold stress against 90 minutes of cold stress (Fig. S4A) for *E. coli* from the literature^56^ showed a very strong correlation (Fig. S4D). Similarly, transcriptomics profile of 40 minutes of oxidative stress against 90 minutes of oxidative stress (Fig. S4B) for *E. coli* from the literature^56^ also showed a very strong correlation (Fig. S4D). Thus, like *M. tuberculosis*, for temporal progression of similar stress, *E. coli* may have a similar stress response mechanism. Interestingly, transcriptomics profile of 90 minutes of cold stress against 90 minutes of oxidative stress (Fig. S4C) showed a very strong correlation (Fig. S4D). Thus, under different stress responses, the transcriptional response of *E. coli* is very similar to that of CTL.

For *E. coli*, ppGpp is a global stress regulator^57^. When an uncharged tRNA binds in the ribosome, the ribosome-associated *relA* protein is activated to synthesize ppGpp^58^, which acts as a global regulator of transcription by modulating transcription complexes at promoters^57^. The stringent response serves to stop the synthesis of stable RNA species, such as rRNA and tRNA, to increase protein degradation pathways to maintain growth rate^58^. Collectively, these responses serve to overcome the starvation. *SpoT* is a cytosolic bifunctional enzyme with ppGpp synthase and hydrolase activity that helps control the levels of ppGpp. CTL does not have homologues of *relA* and *spoT* and does not synthesize ppGpp^59^. The loss of these genes has likely occurred through reductive evolution as a means for adapting to obligate intracellular environment. Thus, highly correlated gene expression under different stress conditions, the absence of *relA* and *spoT* homologues, the inability to synthesize ppGpp, and the upregulation of stable RNA^12^ indicate CTL does not engage in a stringent response during starvation, manifesting as an overlapping transcriptional response during iron and tryptophan starvation.

To further scrutinize the effect of ppGpp for CTL, we reconstructed a model of CTL, capable of producing ppGpp, by adding *spoT, relA, ppx*, and *gpp* proteins to the *i*CTL278. This was performed by adding three reactions to the *i*CTL278 as shown in Fig 4. L. CTL *dksA* does not have the binding pocket for ppGpp. Thus, we also assumed that *dksA* now has a binding site for ppGpp. With the capability to produce ppGpp, it was expected to see an increased biomass growth rate in all tested stress conditions compared to the strain with no capacity to produce ppGpp. Indeed, relative biomass growth rates for all tested stress conditions increased significantly after adding the *spoT* and *relA* proteins to *i*CTL278 (Fig 4. L), compared to the original conditions (4% to 8%). This is a clear indication that ppGpp produces a stringent action under stress conditions by maintaining the biomass growth rates. As CTL lacks the capability to produce ppGpp, thus the nature of its response to nutrient stress is not adaptation, but rather induction of abnormal growth.

### Impact of global stress response on *Chlamydia trachomatis* metabolism

We next sought to determine if this lack of stringent action translated to novel insights into the metabolic landscape of tryptophan- or iron-starved CTL. To answer that, we contextualized *i*CTL278 for UTD24, TRP16, TRP24, BPD16, and BPD24 using the E-flux algorithm.

To confirm that the results from contextualized models are merely not the artifact of transcriptomics data, we calculated the correlation matrix between transcriptomics data predicted flux (more information regarding calculating transcriptomics data predicted flux can be found in the E-flux algorithm sub-section of Materials and Method section) and model predicted flux and found very weak correlation between them (Fig. S5 A). We also found that model predicted flux distribution for different stress conditions are highly correlated (Fig. S5 B). This supports the previously found high correlation among different stress conditions from the transcriptomics data analysis (Fig. S4 I).

As CTL enters persistence, the metabolic difference between UTD24 and TRP16/BPD16 can give insight into the impact of CTL global transcriptome rewiring on its metabolism. To find out the metabolic differences between UTD24 and TRP16, we took the TRP16 model and implemented metabolic bottleneck analysis (MBA)^39^. MBA revealed that, for TRP16, allowing phosphoglycerate mutase (*pgm; ctl0091*) from the glycolysis pathway to carry a similar reaction flux as UTD24 resulted in the same biomass growth rates for UTD24 and TRP16 conditions. The resulting reaction fluxes of other reactions of UTD24 and TRP16 were also found to be similar. This result indicates that potential regulation of *pgm* is the metabolic trait of rewiring of the CTL global transcriptomics response as it enters persistence (Fig. 5 A, B). The *pgm* and the upstream reaction phosphoglycerate kinase (*pgk*) showed opposite transcript profiles under TRP16 condition, with the former being increased (Fig. 5 C, D). Similar results were obtained for BPD16. However, increased level of transcripts may not necessarily translate into more proteins^6,12^.

**Fig. 5.**
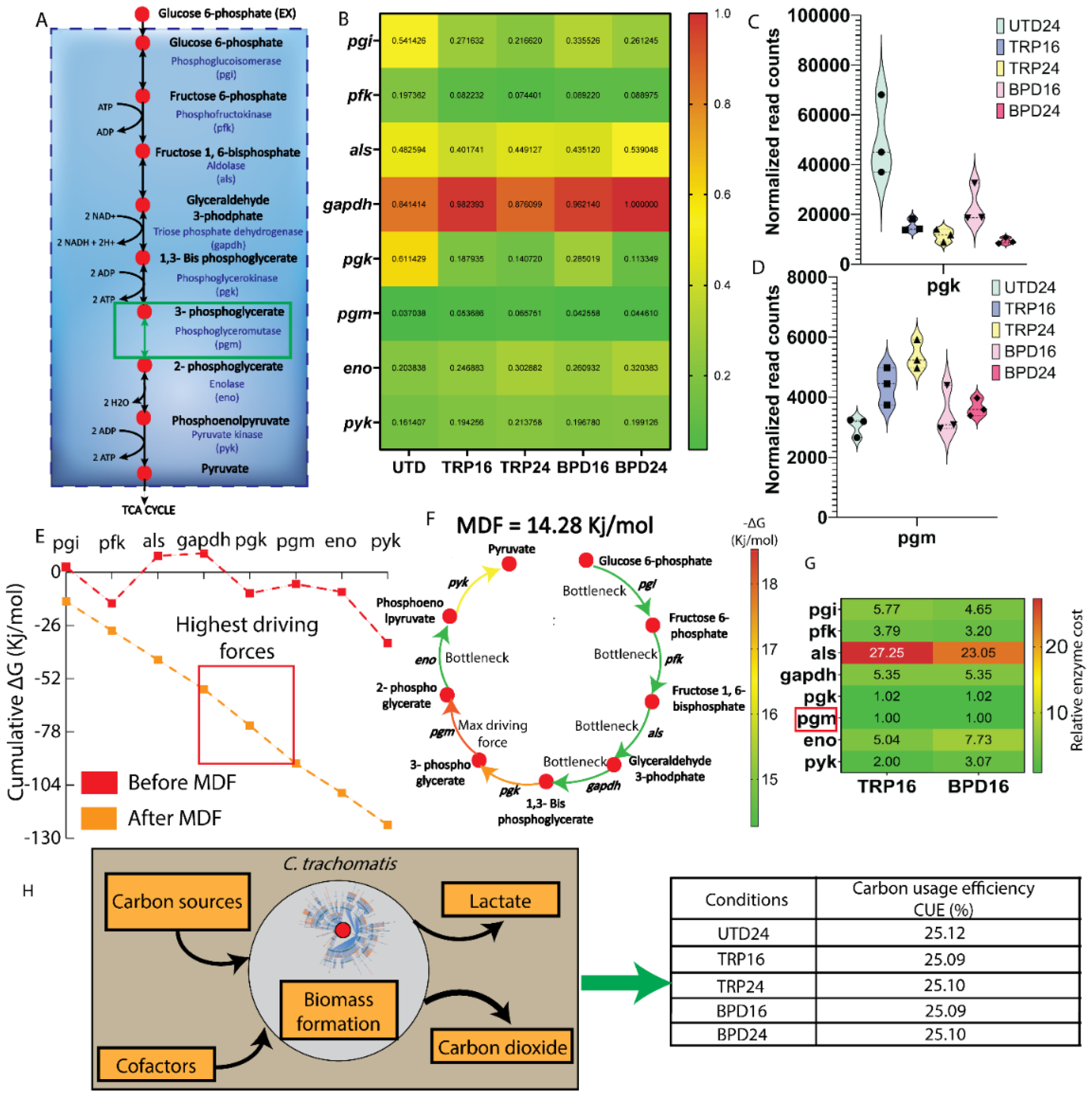
Regulatory reactions in CTL to enter persistence. A) Phosphoglycerate mutase in glycolysis pathway; B) Normalized gene expression heatmap of genes involved in the glycolysis; C) Transcripts level of *pgk* in TRP16-UTD24 and BPD16-UTD24; D) Transcripts level of *pgm* in TRP16-UTD24 and BPD16-UTD24; E) MDF analysis showing the cumulative driving force of glycolysis; E) Driving force plot showing phosphoglycerate mutase (*pgm*) and phosphoglycerate kinase (*pgk*) are two top most driving force reactions; F) Individual driving force of each reactions; G) Enzyme cost of glycolysis for TRP16 showing phosphoglycerate mutase has the lowest enzyme costs; H) Carbon usage efficiency for UTD24, TRP16, TRP24, BPD16, and BPD24.

To dissect further into the significance of *pgm* transcriptional regulation in persistence, we performed max/min driving force (MDF) analysis^60^ on the glycolysis pathway of CTL. MDF analysis maximizes the total driving force of a given pathway within the biologically relevant concentration of different metabolites. Fig. 5 E indicates the driving force of glycolysis before and after the MDF analysis. Among all the reactions in glycolysis, MDF analysis predicted that *pgm* has the highest driving force (Fig. 5 F), while *pgk* has the second-highest driving force (Fig. 5 F). Concentrations of different metabolites, predicted from MDF analysis, are shown in Fig. S5 C. In the context of MDF, shadow price^61^ of a reaction indicates the impact of small perturbation of Gibb’s free energy of that reaction on the overall pathway thermodynamics. Similarly, shadow price of concentrations for each metabolites indicates the impact of small perturbation of metabolite concentration on the overall pathway thermodynamics. Therefore, we calculated the shadow price of driving forces of each of the reaction (Fig. S5 D) and found the first three steps of the glycolysis (*pgi, pfk*, and *als*) having the most impact on the overall driving force with changing reaction fluxes. Next, we calculated the shadow price of concentrations (Fig. S5 E) of each of the metabolites and found H+, glyceraldehyde 3-phosphate, and glycerone phosphate having the most impact on the overall driving force with small changes in concentrations. Flux-force efficacy relationship (Fig. S5 F) indicated a high proportion of each reaction in the forward direction and thus the glycolysis pathway was enzymatically highly efficient. In addition, we calculated the enzyme cost^62^ of each reaction from their enzyme turnover rate, *k*_cat_ (supplementary S3 Table). The turnover rates were calculated using DLKcat for CTL^63^. From the enzyme cost analysis (Fig 5. G), *pgm* has the lowest enzyme costs compared to the other reactions of glycolysis and *pgk* has the second lowest enzyme costs, indicating a low carbon investment to catalyze those reactions compared to other glycolysis reactions. To ensure the observed thermodynamics driving force and enzyme cost implications are not an artifact of reduced carbon usage efficiency (CUE) of CTL, we calculated CUE for all the conditions^64^ and found that CUE remained 25% for both unstressed and stressed conditions (Fig 5. H). Therefore, it is evident that the thermodynamics and enzyme cost of *pgm* could potentially dictate its regulatory role in CTL persistence, rather than its CUE.

### Validation of thermodynamics and enzyme cost analysis through CRISPRi-based suppression and starvation experiment

Using systems biology approaches, we found that *pgm* was regulated at the levels of transcription (from the MBA), thermodynamics driving force, and enzyme cost. However, other glycolysis enzymes, including *pgk*, were regulated only at the level of thermodynamic driving cost and enzyme cost. Therefore, we predicted a *pgm*-mutant would have greater impact on chlamydial growth compared to a *pgk*-mutant, as compared to the wild type.

To test these predictions, *C. trachomatis* inducible knock-down transformants targeting *pgk* or *pgm* were created using CRISPRi as previously described^14^. In these transformants, the guide RNA (gRNA) that targets the promoter region of either *pgk* or *pgm* is constitutively expressed, while the expression of an inactive Cas9 endonuclease (dCas9) is induced with anhydrotetracycline (aTc). The RNA-DNA hybrid formed by the gRNA and genomic DNA is recognized by dCas9, where it binds and blocks RNA polymerase processivity. A control *C. trachomatis* was created by transforming *C. trachomatis* with the L9CRia plasmid that lacks gRNA. The knockdown strains and the empty-vector control were validated for dCas9 induction by aTc, reduction in expression of the target genes, chlamydial development, and bacterial replication (Fig. S6 A, B, and C). In the absence of induction under normal growth conditions, all strains grew equally well (Fig. 6 A).

**Fig. 6.**
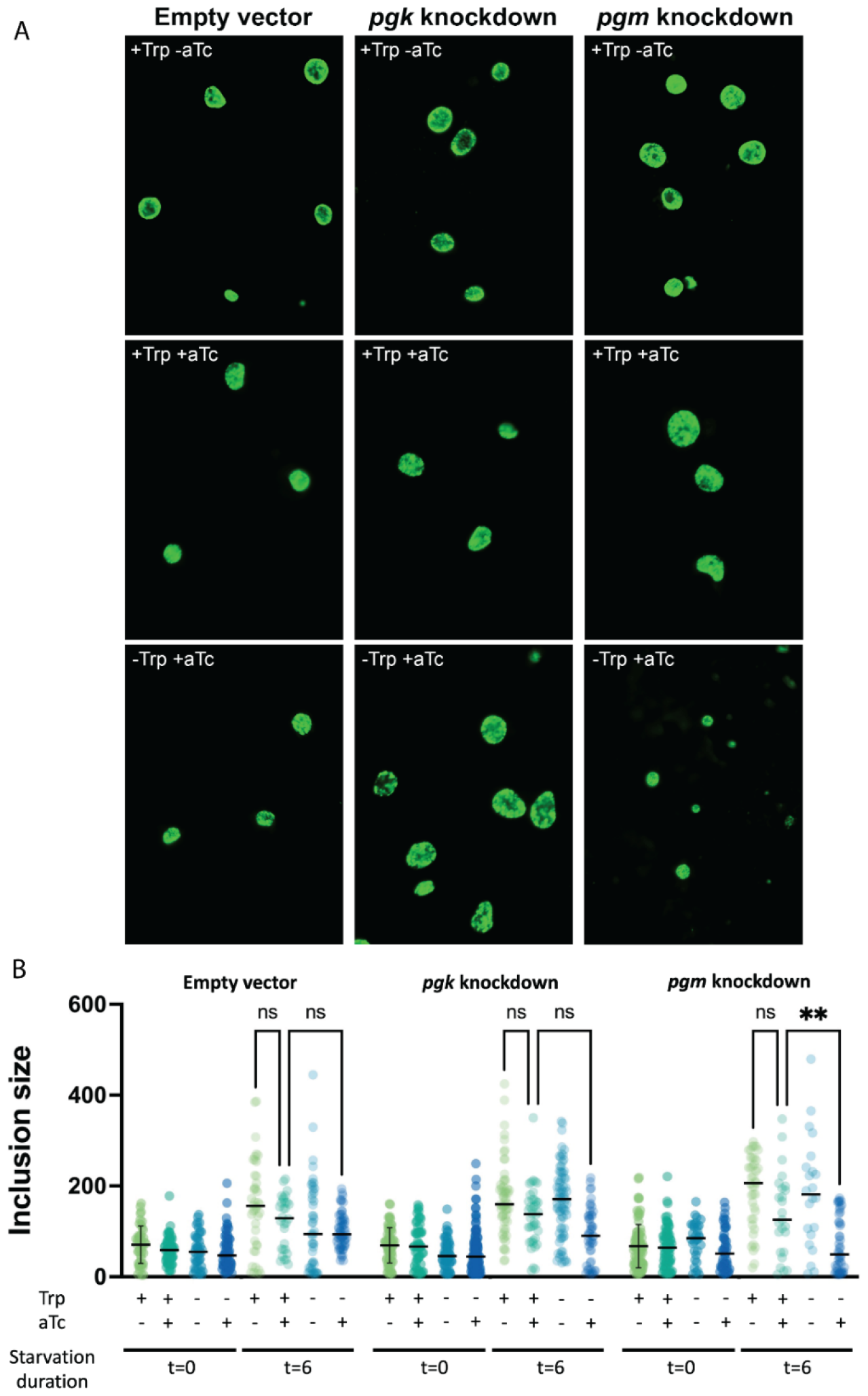
*In vitro* experiments support model prediction that *pgm* plays a major role in pushing *C. trachomatis* to persistence. (A) Tryptophan deprivation for 6 hours resulted in a small reduction in inclusion size. However, with pgm knockdown, the decline became more severe in tryptophan-limited and knockdown-inducing settings. (B) Inclusion reduction became more evident in tryptophan-limited and knockdown-inducing settings, with the pgm knockdown showing a statistically significant difference in inclusion size, corroborating the modeling studies’ findings.

Computational modeling indicated the importance of *pgm* in chlamydial entry into persistence. To validate this prediction, the strains were grown in HeLa cells for 10 h, followed by a 4-h induction of knockdown by treatment with 5 nM aTc. The experimental groups were further subjected to either mock- or tryptophan-starvation for an additional 6 h, with aTc maintained in the growth media. At the end of the experiment (22 h post-infection), samples were fixed and processed for immunofluorescence staining of the chlamydial inclusions. Images were collected by confocal microscopy, and inclusion size measured by NIH ImageJ particle analysis plug-in. Inclusion size correlates with bacterial growth and replication, in that increase inclusion volume is necessary to accommodate increase in bacterial numbers. As shown in Fig. 6 A, all strains yielded similar inclusion sizes in tryptophan-replete media without induction of knockdown (i.e., +Trp, -aTc). Tryptophan starvation for 6 h led to a slight reduction in inclusion size, as expected. However, reduction became more pronounced in tryptophan-limited and knockdown-inducing conditions (i.e., -Trp, +aTc), with the *pgm* knockdown exhibiting statistically significant difference in inclusion size (Fig. 6 B), thus validating the results obtained from the modeling studies. Collectively, these data implicate *pgm* as a critical determinant of sensitivity to tryptophan starvation-mediated persistence.

### Cellular objective of global stress response

As CTL enter the persistence due to the lack of stringent action against the nutrient starvation by regulating *pgm*, it is also critical to investigate the cellular objective of this global transcriptome rewiring of CTL.

To gain insight into the significance of metabolic rewiring associated with tryptophan starvation, we plotted biomass growth rate against different tryptophan uptake rates. If we made more tryptophan available for uptake in the case of TRP24, then it could grow at a higher growth rate compared to TRP16 and UTD24 (Fig. 7 A). A similar result was obtained for BPD16 and BPD24 (Fig. 7 B). We also calculated the correlation matrix of growth patterns (Fig. 7 C and Fig. 7 D) under tryptophan and iron starvation conditions, and like the previous analysis (Fig. 4 I and Fig. S5 A), stress conditions were better correlated.

**Fig. 7.**
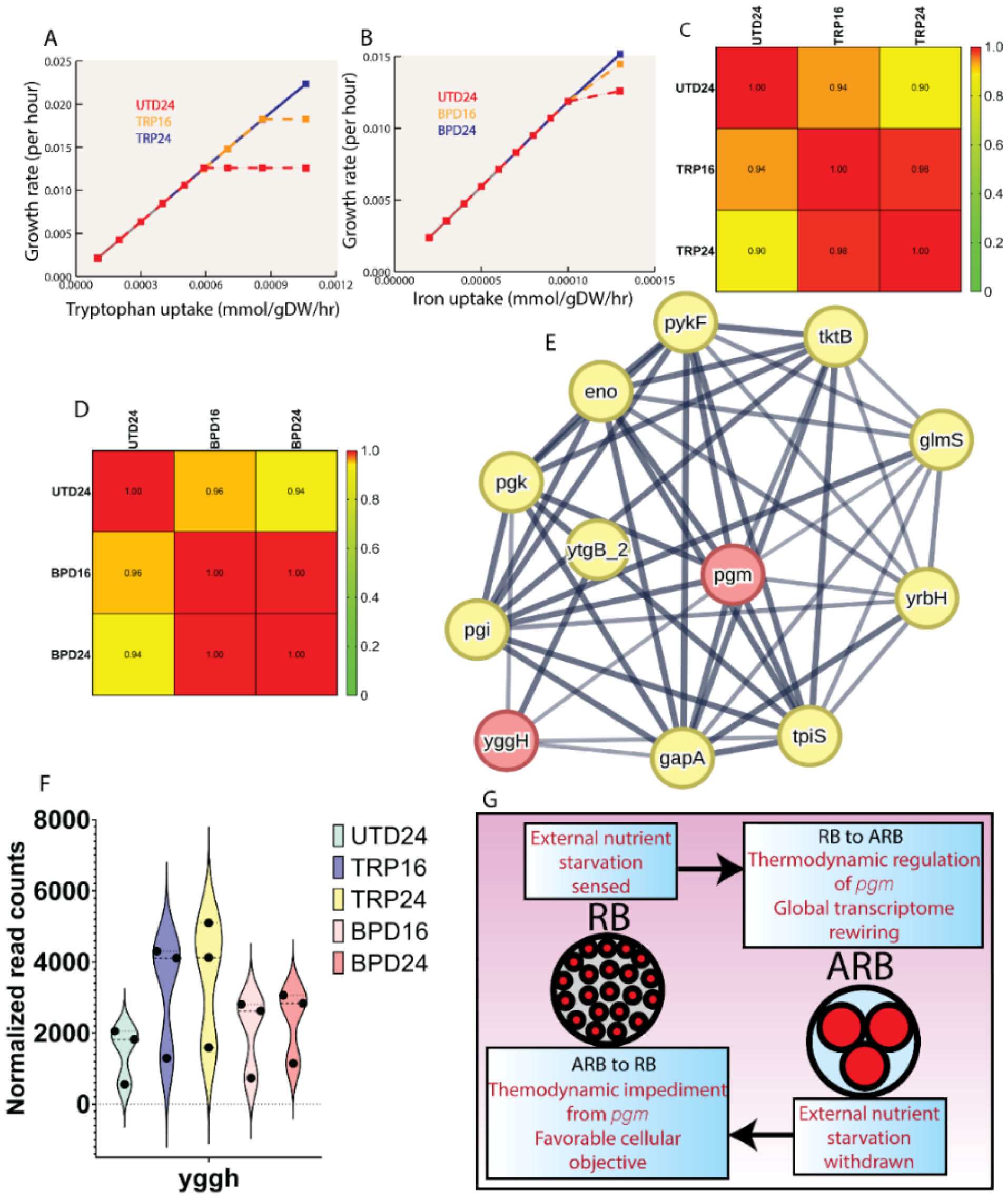
Reaction halting CTL to undergo secondary differentiation to EB. A) The growth rate vs. tryptophan uptake (UTD24, TRP16, TRP24); B) The growth rate vs. iron uptake (UTD24, BPD16, BPD24); C) Correlation matrix calculated among growth rates of UTD24, TRP16, TRP24 using 95% confidence interval and two tail test; D) Correlation matrix calculated among growth rates of UTD24, BPD16, BPD24 using 95% confidence interval and two tail test; E) Protein interaction map for the *pgm* enzyme. Minimum required interaction score was 0.50 and top five interactions allowed for first shell; F) Normalized read counts of *yggH* in different conditions; G) Schematic of the mechanism proposed to enter persistence and factors that keep CTL in the persistence mode.

However, previous study^65^ suggested that, with a shorter starvation timeline, CTL should better restore its growth and start differentiating to the RB upon availability of nutrients. In other words, it should be easier to restore the growth of CTL for 16 hours of nutrient starvation compared to 24 hours of nutrient starvation. Thus, we explored the model further to investigate this phenomenon. We again applied MBA in the TRP16 metabolic model. Surprisingly, we found that if *pgm* reaction flux was relaxed to its value obtained from TRP24 model (Fig. 5 A), the biomass of TRP16 and TRP24 became the same. A similar result was obtained for the BPD16 and BPD24. This analysis indicates that the cellular objective of the CTL global transcriptome rewiring under the persistence state is to reach such a cellular phenotype so that, when the stress is withdrawn, CTL can exit persistence immediately and re-enter the normal development cycle. CTL can do that by relaxing its regulation of *pgm*, which is critical. This final objective also supports the evolutionary selection of CTL, which is to maximize its own fitness function. The protein interaction network suggests that *pgm* has a strong interaction with *yggH* (Fig. 7 E). *YggH* catalyzes the *S*-adenosyl-l-methionine-dependent formation of *N*^7^-methylguanosine at position 46 (m^7^G46) in tRNA and can impact the activity of tRNA^66^. It was previously reported that inhibition of tRNA synthesis can induce persistence in CTL^67^. The normalized read counts of *yggH* shows significantly higher level of transcript in all the stress conditions compared to the UTD24 (Fig. 7 F). Thus, the regulation of *pgm* may be mediated in part by the *yggH*. The proposed mechanism that led CTL to enter persistence is shown in Fig. 7 G. Quantitative proteomics from the literature^68^ indicated that the ARB is primed for a burst in metabolic activity upon nutrient availability, whereas the RB is geared towards nutrient utilization, a rapid increase in cellular mass, and securing the resources for an impending transition into the EB form. In addition to confirming previously published findings, this study indicates that *pgm* may be the possible barrier to ARB-to-RB transition.

In this work, transcriptomics datasets were generated for untreated and nutrient starved CTL. We then used K-mean clustering on those transcriptomics data to identify core and stress-specific components of CTL transcriptome which led to the finding that CTL might have a global transcriptomic rewiring under different stress conditions. This global transcriptomics rewiring along with the absence of ppGpp lead to the lack of stringent action, which is the likely reason of CTL persistence. To further investigate how that global transcriptomic rewiring impacted the metabolism of CTL, we reconstructed its genome-scale model *i*CTL278 and later contextualized it for normal and stress conditions with transcriptomics data. Using MBA on these contextualized *i*CTL278, we proposed *pgm* regulates the entry of CTL to persistence. Later, thermodynamics driving force, enzymatic cost, CRISPRi-based suppression, and subsequent starvation experiments supported this finding. Furthermore, we investigated the cellular objective of the persistence and found that priming itself for a metabolic burst upon availability of nutrient dictates the nature of the persistence. MBA further implicated *pgm* preventing ARB to convert back to the RB. Overall, this study showed how a combined systems and synthetic biology approach can lead to a better understanding of CTL persistence. Future research effort will be geared towards studying the interaction between CTL and different host epithelial cells (e.g., endocervical and endovaginal epithelial cell) and similar to this study, finding the metabolic signatures influencing the CTL infection in these cells.

## Materials And Methods

### Cell lines

Human female cervical epithelial adenocarcinoma HeLa cells (RRID: CVCL_1276) were cultured at 37°C with 5% atmospheric CO_2_ in Dulbecco’s Modified Eagle Medium (DMEM; Gibco, Thermo Fisher Scientific) supplemented with 10□μg/mL gentamicin, 2□mM l-glutamine, and 10% (vol/vol) filter sterilized fetal bovine serum (FBS). For all experiments, HeLa cells were cultured between passage numbers 3 and 15. HeLa cells were originally authenticated by ATCC via STR profiling and isoenzyme analysis per ATCC specifications.

### Bacterial strains

*Chlamydia trachomatis* serovar L2 was originally obtained from Dr. Ted Hackstadt (Rocky Mountain National Laboratory, NIAID). Chlamydial EBs were isolated from infected HeLa cells at 36 to 40 hpi and purified by density gradient centrifugation essentially as described^69^. For infections, at 80 to 90% confluence, HeLa cells were first washed with Hanks Buffered Saline Solution (HBSS; Gibco, Thermo Fisher Scientific) and ice-cold inoculum prepared in HBSS at the indicated multiplicity of infection was overlaid onto the cell monolayer. To synchronize the infection, inoculated cells were then centrifuged for 15□min at 500xRCF, 4°C in an Eppendorf 5810 R tabletop centrifuge with an A-4-81 rotor. The inoculum was then aspirated and pre-warmed DMEM (or relevant media with treatment supplementation) was added to the cells. Infected cultures were then returned to the tissue culture incubator until the indicated time post-infection.

### Treatment conditions and induction of knockdown of *pgk* and *pgm* transcription by CRISPRi

HeLa cells were infected at a multiplicity of infection (moi) of 0.5 (t=0 hour post-infection (hpi)), and were maintained in complete DMEM for 14 h. The infected cells were then exposed to anhydrous tetracycline (aTc) at 5 nM for 4 h to induce knockdown of expression (t=14 hpi). Tryptophan depletion was performed at t=18 hpi by first washing cells with HBSS and then replacing complete DMEM with tryptophan-depleted DMEM-F12 (U.S. Biological Life Sciences). Treated cells were then returned to the tissue culture incubator for the remainder of the experimental time course. At t=22 hpi, samples were collected and fixed with freshly prepared 4% paraformaldehyde and permeabilized for 10 min with 0.1% Triton X-100 in PBS. Samples were immunostained with mouse antibody against the *C. trachomatis* major outer membrane protein (1:1000 dilution). Samples were incubated at 4C overnight with constant rocking. The next day, the primary antibody solution was removed and samples rinsed 3x with 1x PBS, and incubated with 1:1000 dilution of goat anti-mouse IgG conjugated with Alexa-488 for 1 h. Samples were rinsed and visualized by fluorescence microscopy. Images were processed and inclusion size measured using NIH ImageJ.

### Genome-scale metabolic model reconstruction of CTL

The genome-scale metabolic network reconstruction was based on available genome information of the model strain *C. trachomatis* L2, according to the ModelSEED databases^70^, BRENDA^71^, and available literature. Additional reactions were added to the model based on the previously published genome-scale metabolic model of CTL^29^, experimental data^8^, and previously published literature on metabolic traits of CTL. Gap filling for pathways in our model were first conducted in Kbase^40^ and then by OptFill^42^ based on the complete media, as complete media was used in the experimental settings as well to grow CTL *in vitro*. In total, 95 reactions were added to the draft model to fill the gaps. The model was further checked for elemental mass balance and MEMOTE reports confirms that all the reactions in the model are mass balanced. GPR for all the reactions were manually curated from the KEGG database. As a result, 525 out of 692 reactions has GPR relationship. We used GAMS platform along with CPLEX solver for solving all the optimization problems. NEOS server can be used to run GAMS codes without having to buy the license. Details procedure of running GAMS codes in NEOS server can be found in the literature^72^. However, for the convenience of COBRApy users, SBML version of the *i*CTL278 is also provided.

### Flux sampling and tSNE plot of *i*CTL278

Flux sampling was performed using the ACHR sampling toll of COBRApy. For tSNE, plot we used a perplexity of 45 and random state of 42. Clusters from the tSNE plot was generated using K-mean clustering algorithm.

### Parsimonious flux balance analysis

pFBA^20^ is constrained based optimization technique to model GSMs. The pseudo-steady state mass balance in pFBA is represented by a stoichiometric matrix, where the columns represent metabolites, and the rows represent reactions. For each reaction, upper and lower bounds is imposed based on thermodynamic information. pFBA provides the flux value for each reaction in the model according by solving the following optimization problem:

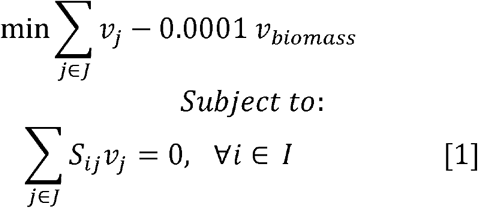

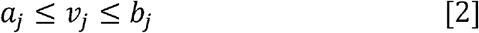

In this formulation, *l* is the set of metabolites and *J* is the set of reactions in the model. *S*_*ij*_ is the stoichiometric matrix with i indicating metabolites and *J* indicating reactions, and *v*_j_ is the flux value of each reaction. The objective function, *v*_*biomass*_, is the proxy of the growth rate of an individual cell. *a*_j_ and *b*_j_ are the lower and upper bounds of flux values for each reaction. For forward reactions, the highest possible bounds were 0 mmol/gDW/h to 1000 mmol/gDW/h. For the reversible reactions, the highest possible bounds were -1000 mmol/gDW/h to 1000 mmol/gDW/h.

### Flux variability analysis

Result from FBA may include degenerate optimal solutions, this flux variability analysis (FVA)^19^ was used to find out the alternate flux distributions. The formulation is the following:

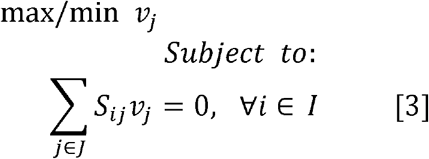

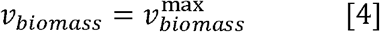

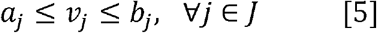

FVA maximizes and minimizes each of the reaction fluxes subject to the pseudo-steady mass balance, fixing biomass growth rate for a specific condition, and upper and lower bounds on reaction fluxes. In this manuscript, all the FVA analysis resulted in very tight bounds (changes can only be observed after two decimal place).

### E-Flux algorithm

E-Flux is an extension of FBA/pFBA that uses transcriptomic data to further constrain the feasible space based on the transcriptomics data^37^. The E-flux algorithm involved solving the following linear optimization problem:

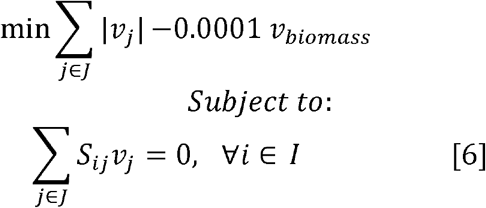

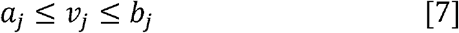

where *a*_j_ and *b*_j_ are the minimum and maximum allowed fluxes through reaction *J*, based on the transcriptomics data. The E-Flux method calculates the upped bound, *b*_j_, for the *j*^th^reaction according to the following function of the gene expression:

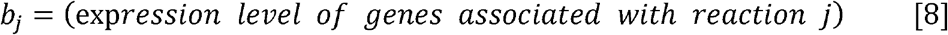

In this manuscript, *b*_j_ is the exact level of each reaction that was calculated through its GPR association (i.e., based on “OR” (addition of gene expressions), and “AND” (minimum of gene expressions) relation). If the reaction catalyzed by the corresponding enzyme was reversible then *a*_j-_= *-b*_j_, otherwise *a*_j_ =0.

### Flux sum analysis

Metabolite pool size of ATP in the validation section was determined based on the flux-sum analysis (FSA) method^32^. The flux-sum is a measure of the amount of flux through the reactions associated with either the production or consumption of the metabolite. The range of the flux-sum can be calculated as follows as follows:

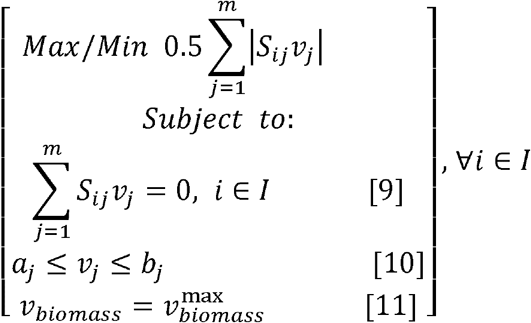

Here, set *l* represents the set of metabolites for which flux sum will be calculated. By linearizing the objective function, the resulting formulation became a mixed-integer linear programming problem. Therefore, the basic idea was to determine the range of the flux-sum of a metabolite under a given condition by fixing the maximum biomass growth of that condition (equation 11).

### K-mean clustering

K-mean clustering algorithm was used to classify different genes into different clusters. Number of clusters was determined using the Elbow method (Fig. S7). The whole K-mean clustering was implemented in Python, using numpy, pandas, and sklearn modules. Different plots for the K-mean clustering was generated using matplotlib module. Default setting of K-mean clustering, mentioned in the sklearn, was not changed in this study.

### Metabolic bottleneck analysis

To determine the metabolic bottleneck in a GSM metabolic bottleneck analysis^39^ was used. The formulation is as following.

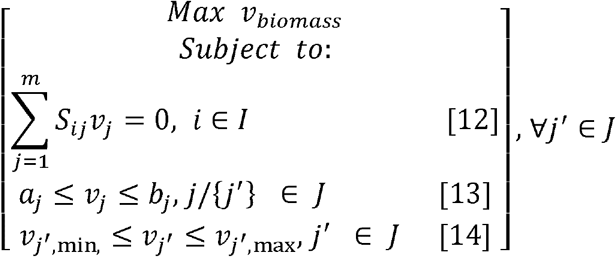

Here *a*_j_ is the lower bound reaction *v*_j_ and *b*_j_ is the upper bound of reaction *v*_j_ . Both *a*_j_ and *b*_j_ were calculated from the transcriptomics data and gene-protein-reaction association. *v*_j_^′^_,min_ is the expanded lower bound of the reaction *J* ^′^and *v*_j_^′^_,max_ is the expanded upper bound of the reaction*J*^′^. In this case, we set 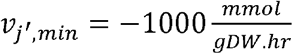 and 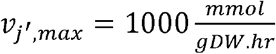. We solved the optimization problem by maximizing the biomass *v*_*biomass*_ for the new expanded flux space of each reaction *J*^′^in an iterative manner and then recorded the biomass growth rate. From this biomass growth rate collections, we can check for which *J*^′^ biomass growth rate increased significantly. Then that*J*^′^can be considered as the metabolic bottleneck of a given metabolic network.

### Max/Min driving force analysis

To find out the thermodynamic driving force and thermodynamic bottleneck of a given pathway, we used max/min driving force analysis (MDF) from the literature^60^. In this method, we maximized the driving force of each reaction in a given pathway within the biologically relevant concentration and found the maximum possible driving force of a pathway. The formulation of MDF analysis is given below:

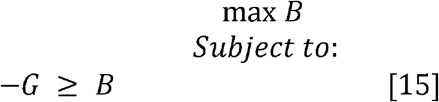

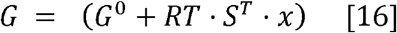

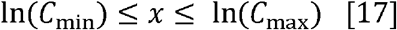

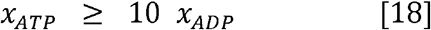

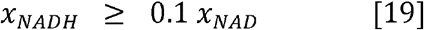

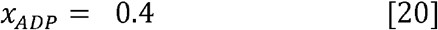

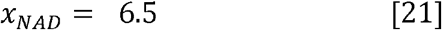

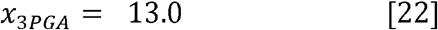

*G*^0^ is the standard Gibbs free energy, *R* is the gas constant, *T* is the temperature, and *x and C* indicates concentration. This analysis is particularly useful when metabolomics information is not fully available for a given pathway. Thus biologically relevant metabolite concentration range can be used to infer information about a pathway, whether it will be thermodynamically feasible or not. Furthermore, this analysis will indicate reaction(s) with the highest and lowest thermodynamic driving forces and will also provide concentration of metabolites to support the maximum pathway driving force. Here we fixed the ATP/ADP and NADF/NAD ratio as mentioned in the original MDF article^60^. Besides concentrations (mM) of ADP, NAD, and 3-PGA were fixed in the MDF optimization problem based on literature evidence^73,74^. Shadow price of different constraints were calculated using the built-in features of GAMS. The temperature in the optimization problem was set to 37°C, which is similar to the temperature of the growth culture. Ratio of forward to backward reaction fluxes for each of the glycolysis reaction was calculated using the following equation.

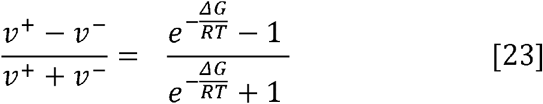

Here, Δ*G* was calculated from the MDF analysis. Also, *v*^+^and *v*^-^ indicate reaction flux in forward and backward directions respectively.

### Enzyme cost calculation

To find out enzyme synthesis costs for different enzymes of a given pathway, we used the following equation from the literature^62^ assuming all the enzymes are fully saturated.

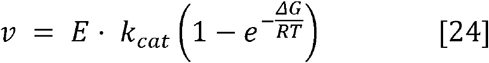

Here, *v* is the reaction flux obtained from the contextualized GSM, *E* is the enzyme cost, *k*_cat_ is the enzyme turnover rate, Δ*G* is the gibbs free energy obtained from the MDF analysis, *R* is the gas constant, and *T* is the temperature. Relative enzyme cost was calculated using the following equation.

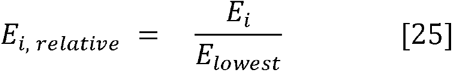

Here, *E*_*i,relative*_ is the relative cost of an enzyme in a given pathway, *E*_i_ is the actual cost of an enzyme in a given pathway, and *E*_*lowest*_ is the lowest cost of an enzyme in a given pathway.

### Simulation platform

The General Algebraic Modeling System (GAMS) version 24.7.4 with IBM CPLEX solver was used to run pFBA and FVA, E-Flux, FSA, MBA, and MDF algorithm on the model. Each of the algorithm was scripted in GAMS and then run on a Linux-based high-performance cluster computing system at the University of Nebraska-Lincoln. Furthermore, flux sampling, tSNE plot, and K-mean clustering analysis of transcriptomics data was performed in Python using an Intel(R) Core(TM) i5-8250U CPU @ 1.60GHz HP laptop with 8.00 GB of RAM and 64-bit operation with Windows 11 Home operating system.

## Supporting Information

S1 Text. MEMOTE report of *i*CTL278.

S1-S7 Figure. Supporting figures.

S1 Table. Details about clusters from flux sampling.

S2 Table. Details about each clusters.

S3 Table. Details of *k*_*cat*_ for each reaction in glycolysis.

## Acknowledgments

RS gratefully acknowledge the funding support from National Science Foundation (NSF) CAREER grant (1943310), National Institute of Health (NIH) R35 MIRA grant (5R35GM143009), and Nebraska Collaboration Initiative grant. RC, SO, and EAR acknowledge AI132406 from the NIH.

## Author Contribution

Conceptualization: Rajib Saha, Rey A. Carabeo

Data curation: Niaz Bahar Chowdhury

Formal analysis: Niaz Bahar Chowdhury

Funding acquisition: Rajib Saha, Rey A. Carabeo, Elizabeth A. Rucks, Scot P. Ouellette

Methodology: Niaz Bahar Chowdhury, Nick Pokorzynski

Project administration: Rajib Saha, Rey A. Carabeo, Elizabeth A. Rucks, Scot P. Ouellette

Software: Niaz Bahar Chowdhury

Supervision: Rajib Saha, Rey A. Carabeo

Validation: Niaz Bahar Chowdhury

Visualization: Niaz Bahar Chowdhury

Writing – original draft: Niaz Bahar Chowdhury

Writing – review & editing: Rajib Saha, Rey A. Carabeo

## Data Availability

The data that support for the findings of this study can be found in the related cited articles and/or in the supplementary data. All the codes used to generate these results can be accessed in the GitHub repository (https://github.com/ssbio/c_trachomatis).

## Notes

### Competing Interest Statement

The authors have declared no competing interest.

